# Large-scale in-cell photocrosslinking at single residue resolution reveals the molecular basis for glucocorticoid receptor regulation by immunophilins

**DOI:** 10.1101/2023.01.16.524346

**Authors:** Asat Baischew, Sarah Engel, Thomas M. Geiger, Felix Hausch

**Affiliations:** Department of Chemistry, Technical University Darmstadt, Darmstadt, Germany

## Abstract

The large immunophilins FKBP51 and FKBP52 play key roles in the Hsp90-mediated maturation of steroid hormone receptors, which is crucial for stress-related disorders and correct sexual embryonic development, respectively ^1–3^. A prominent regulatory target is the glucocorticoid receptor (GR), whose activation is repressed by FKBP51 ^4,5^ and facilitated by FKBP52 ^6,7^. Despite their vital roles, the molecular modes of action of FKBP51 and FKBP52 are poorly understood since the transient key states of FKBP-mediated GR-regulation have remained experimentally elusive. Here we present the architecture and functional annotation of FKBP51-, FKBP52- and p23-containing Hsp90-apoGR preactivation complexes, trapped by systematic incorporation of photoreactive amino acids ^8,9^ inside human cells. The identified crosslinking sites depended on a functional Hsp90 chaperone cycle, were disrupted by GR activation, and clustered in characteristic patterns, defining the relative orientation and contact surfaces within the FKBP/p23-apoGR complexes. Strikingly, GR binding to the FKBP^FK1^ but not the FKBP^FK2^ domains were modulated by FKBP ligands, explaining the lack of FKBP51-mediated GR derepression by certain classes of FKBP ligands. These findings show how FKBP51 and FKBP52 differentially interact with the apoGR ligand binding domain, they explain the differentiated pharmacology of FKBP51 ligands, and provide a structural basis for the development of FKBP ligands with higher efficacy.

---

Steroid hormone receptors are key endocrine effectors that rely on the Hsp90 chaperone machinery ^10,11^, assisted by a various co-chaperones. Among these, FK506-binding Proteins 51 and 52 (FKBP51 and FKBP52) are thought to fine-tune the final steps of steroid hormone receptor maturation. Their physiological importance has become evident by transgenic studies ^12^, where FKBP52 reduction or deficiency severely compromised steroid hormone signaling in cells ^13,14^ and sexual embryonic development and glucose homeostasis in mice ^12^. Conversely, FKBP51 knockout mice were protected from diet-induced obesity ^4,15^ or various forms of chronic pain ^16^ and displayed an enhanced stress coping behavior ^17,18^. FKBP51 expression is robustly induced by steroid hormones and various types of stress in cells, mice and humans, further underscoring the prominent role of FKBP51 as a key regulator of steroid hormone receptors and stress physiology ^19^. The importance of FKBP51 (encoded by the *fkbp5* gene) for human health is supported by FKBP51-hyperinducing single nucleotide polymorphisms that have repeatedly been associated with stress-related disorders ^20^. Collectively, FKBP51 has emerged as a potential target to treat stress-related disorders, obesity-induced diabetes, or chronic pain ^2,21^. Despite substantial recent structural advances on Hsp90-mediated nuclear hormone maturation ^22–26^, a detailed mechanistic understanding of the key steps of GR regulation by the large FKBP co-chaperones has remained elusive, in part due to difficulties to purify or functionally reconstitute FKBP51 and FKBP52 together with GR in defined biochemical systems ^27,28^. Here we present a detailed molecular description of FKBP51- and FKBP52-containing Hsp90-GR complexes in an authentic cellular environment, that captures GR in the state of regulation by FKBP51 and FKBP52 immediately prior to activation.

## FKBP51 and FKBP52 have large and defined GR interaction interfaces

To interrogate the architecture of the FKBP51/52-Hsp90-GR heterocomplexes in mammalian cells (HEK293), we resorted to amber suppression ^8,9^ to site-specifically incorporate the photoreactive unnatural amino acid para-benzoyl phenylamine (pBpa) at defined positions in different members of the complexes. This was followed by life cell irradiation and analysis of photocrosslinking adducts.

As a technical proof-of-concept, we first investigated amino acid positions in the TPR domain of FKBP51 (FKBP51^TPR^), which are predicted to mediate key contacts to the C-terminal domain of Hsp90 (Hsp90^CTD^) ^29^. Indeed, several positions in FKBP51^TPR^ were identified, which upon UV irradiation produced higher-molecular weight complexes corresponding to crosslinked FKBP51-Hsp90 heterodimers (Extended Data Fig. 1a). This included positions previously shown to be in tight contact with Hsp90 (Extended Data Fig. 1d and e) ^29^ as well as positions poised to interact with the unresolved linker proceeding the C-terminal MEEVD motif of Hsp90 (Hsp90^MEEVD^, Extended Data Fig. 1b).

We then set out to map the interaction interface of FKBP51 with GR, individually incorporating pBpa into all surface-exposed amino acid positions of FKBP51 ^30^. Out of the 216 tested sites, 46 crosslinks of FKBP51 to the GR were detected by Western Blot (exemplarily shown in Fig. 1a and fully shown in Extended Data Fig. 2a). These segregated into well-defined interaction sites spanning all three domains of FKBP51. The GR-reactive amino acid positions clustered around the FK506-binding site (e.g., R73, E75, Q85, P120) of the FK1 domain, the β1-β2 loop (e.g., Y159), the tip of helix α1 (e.g., E207) and the α1-β4 loop (e.g., Q210) of the FK2 domain, the FK2-TPR linker (e.g., W257), and the α2-α3 loop (e.g., E301) and helix α4 (e.g., K342) of the TPR domain (Fig. 1b and Extended Data Fig. 2c). Mapping of the crosslinking sites on the FKBP51 structure ^30^ revealed a large and continuous interaction surface on adjacent sides of the FK1, FK2 and the tip of the TPR domain (Fig. 1b). Strikingly, not a single crosslink was detected on the opposite side of FKBP51, indicating the presence of a well ordered and reasonably populated FKBP51-Hsp90-GR complex inside human cells.

**Fig. 1.**
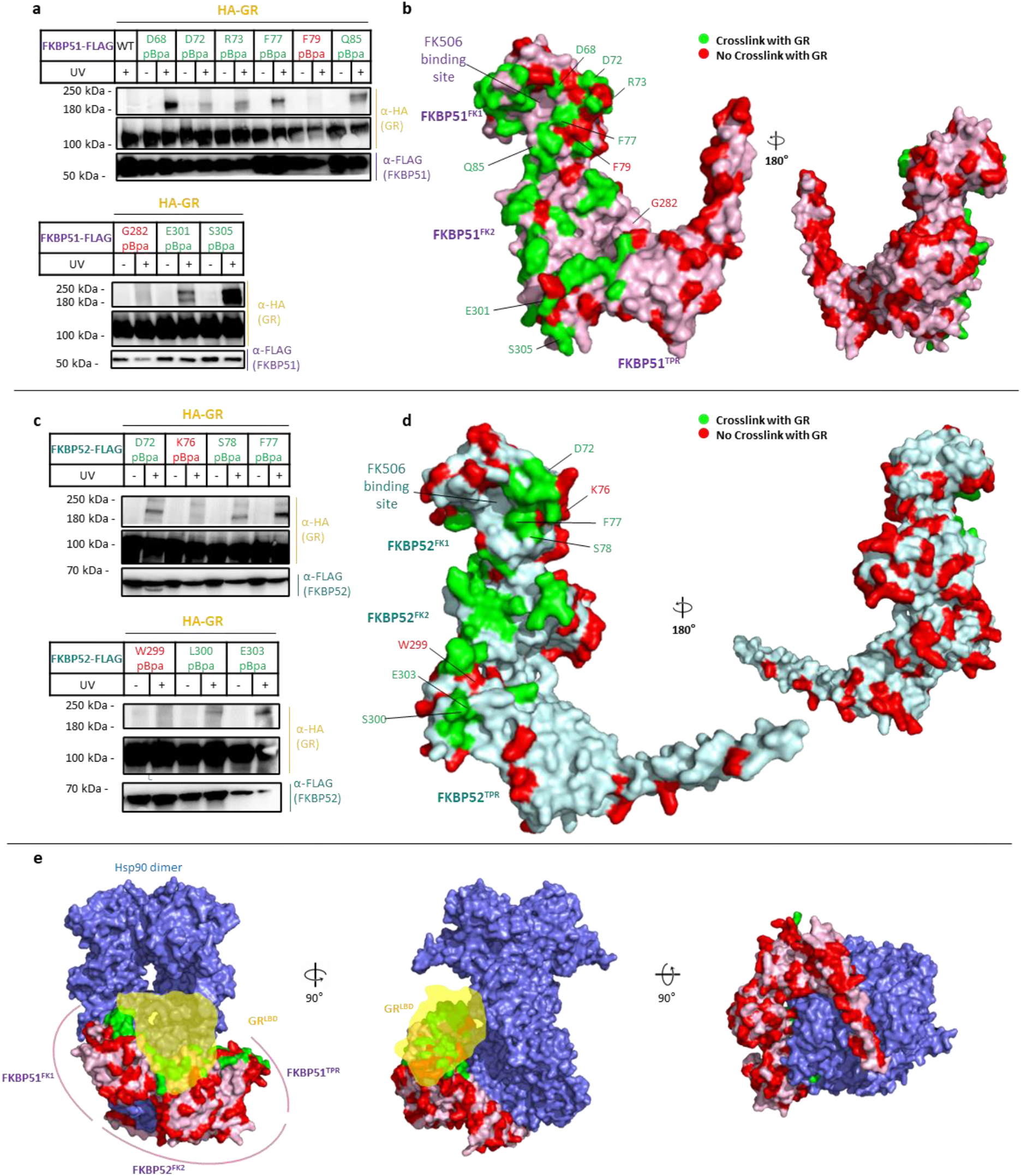
Large-scale in-cell photocrosslinking reveals large and defined interaction interfaces of FKBPs with GR. **a,** Western blots of exemplary FKBP51 pBpa mutants expressed and photocrosslinked in HEK293 cells co-overexpressing HA-tagged GR. UV light-induced HA-reactive bands at a size of approx. 180 kDa are indicative of the mutated position being in proximity to GR. **b,** GR-photoreactive positions (highlighted in green) and inactive position (shown in red) in FKBP51 (pale pink), identified by Western blotting as in **a**, were mapped on the structure of FKBP51 (PDB: 5OMP). **c,** In-cell photocrosslinking analysis for FKBP52, performed identical as in **a**. **d,** FKBP52 crosslinks to GR were mapped on the predicted FKBP52 structure (AFQ02790). **e,** FKBP51→GR crosslinks are mapped on the Hsp90-FKBP51 complex (PDB: 7J7I, p23 omitted for clarity) to illustrate the orientation of the FKBP51-GR interaction interface. The broad position of Hsp90-bound GR^LBD^, estimated from PDB-ID 7KRJ, is indicated as a transparent yellow shape.

Next, we similarly characterized the interface of FKBP52 with GR (Fig. 1c and Extended Data Fig. 2b). Out of a library of 153 pBpa mutants, 29 GR-photoreactive positions between FKBP52 and GR were detected. Again, these clearly segregated on one side of the FK1, FK2 and part of the TPR domains of FKBP52, whereas no crosslinks were detected on the other side (Fig. 1d). To our surprise, the overall GR-crosslinking patterns of FKBP51 and FKBP52 were almost identical (Fig. 1b and d, Extended Data Fig. 2c), revealing a common basal binding mode of these co-chaperones with GR.

The broad orientation of this binding mode became apparent when mapping the crosslinks on the structure of FKBP51 in complex with Hsp90 (Fig. 1e) ^29^. Notably, all GR-reactive positions on FKBP51 are oriented towards the GR binding site of Hsp90 that was suggested from recent cryo-EM structures of GR-Hsp90 complexes ^22,23^. While it has been clear from these structures that GR and/or FKBP51 have to rearrange in a FKBP51-Hsp90-GR complex, the details of this rearrangement have been unclear. Our data support a model where minor movements of FKBP51 or FKBP52 away from the core of Hsp90 are sufficient to functionally interact with GR and represent the predominant conformation of FKBP51 and FKBP52 in FKBP-GR-containing complexes inside cells.

## FKBP51 wraps around the GR ligand binding domain inside cells

To further refine the orientation of GR in the FKBP51-Hsp90-GR complex we generated ^31^ and screened a pBpa mutant library (215 mutants) covering the putative surface of full-length GR, including 42 surface-exposed amino acid positions in the GR ligand binding domain (GR^LBD^), 24 residues in the DNA-binding domain, and 149 residues in the structurally uncharacterized N-terminal part of GR. Several FKBP51-reactive crosslinks were identified in the GR^LBD^ (Fig. 2a), while none were observed in the N-terminal parts (Extended Data Fig. 3), confirming the GR^LBD^ as the relevant interaction domain for FKBP51 in cells. Within the GR^LBD^, one interaction hotspot centered around helix α12 (e.g., T744, E755, and N768, Fig. 2b and Extended Data Fig. 4a and b). Mapping the crosslinks on the structure of the GR^LBD^-Hsp90 complex ^23^ revealed another crosslinking hot spot on the other side of the GR^LBD^ (e.g., R614, R655, and S659, Fig. 2b), which are located at the tip of helix α5 and in the α7-α8 loop. These findings, especially for N768 as the strongest GR→FKBP51 crosslink observed, require the GR^LBD^ to rearrange compared to the orientation observed in the structure of the Hsp90-GR complex. We propose a clockwise rotation (viewed from Fig. 2b left) to satisfy all crosslinks observed. The full set of GR→FKBP51 crosslinks reveals a complex, where FKBP51 wraps around the GR^LBD^, specifically from helix α12 around helix α3 and the α1-α3 loop to reach the tip of helix α5 and the α7-α8 loop (Extended Data Fig. 4b).

**Fig. 2.**
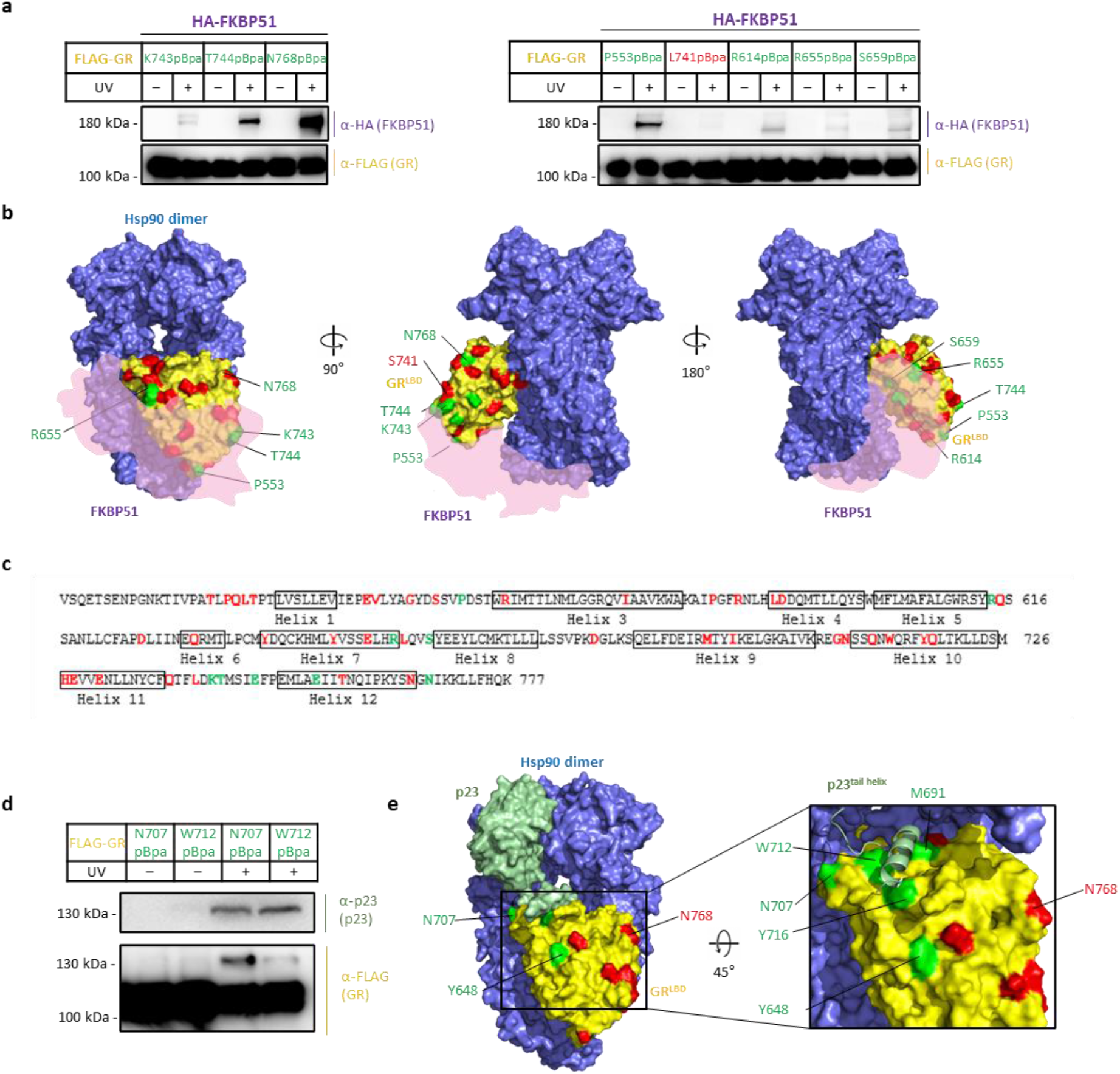
Large scale photocrosslinking of the full-length GR in mammalian cells confirms the interaction of the GR ligand binding domain and co-chaperones FKBP51 and p23. **a,** Western blots of exemplary full-length GR pBpa mutants expressed and photocrosslinked in HEK293 cells co-overexpressing HA-tagged FKBP51. UV light-induced HA-reactive bands at a size of approx. 180 kDa are indicative of the mutated position being in proximity to FKBP51. **b,** GR→FKBP51 crosslinks were mapped on the structure of the Hsp90-GR complex (PDB: 7KRJ, p23 omitted for clarity) with crosslinks highlighted in green and inactive position indicated in red. The broad position of Hsp90-bound FKBP51, estimated from PDB-ID 7L7I, is indicated as a transparent light pink shape. **c,** Crosslinks or inactive positions are mapped to the sequence of GR^LBD^ (523–777), with indication of secondary structure. **d,** Western blots of exemplary full-length GR pBpa mutants expressed and photocrosslinked in HEK293 cells (without FKBP51 co-overexpression). UV light-induced FLAG-reactive bands and p23-reactive bands at a size of 130 kDa are indicative of the mutated position being in proximity to p23. **e,** GR→p23 crosslinks were mapped on the structure of the Hsp90-GR-p23 complex (PDB: 7KRJ) with crosslinks highlighted in green and inactive position indicated in red. In the close-up view the p23tail helix (p23 120-130) is shown as cartoon.

## Exploration of p23-GR contacts

With the GR single point mutant library in hand, we also explored crosslinks of GR with the Hsp90 co-chaperone p23, which is reported to be essential for GR maturation ^32^. Numerous GR→p23 crosslinks were observed that clustered on the GR (Fig. 2d and Extended Fig. 4c), e.g., at positions Y648, R655, M691, I694, N707, W711, and Y716). This pattern fits remarkably well to a recent cryo-EM structure of the p23-Hsp90-GR^LBD^ complex ^23^ (Figure 2e). Some GR^LBD^→FKBP51 crosslinks (R614, R655 and S659) (Fig. 2b) are suspiciously close or even identical to GR^LBD^→p23 interaction sites. This suggests that FKBP51 may compensate for some of the GR-p23 contacts lost due to the GR rearrangement postulated above.

## Both FKBPs and p23 interact with apo-GR^LBD^ in an Hsp90-dependent manner

To further define the functional characteristics of the FKBP51-, FKBP52-, and p23-GR-Hsp90 complexes in cells we used a selected set crosslinking positions as intracellular proximity sensors. Treatment of cells expressing pBpa-incorporating FKBP51 mutants (at D68, E75 or Q210) with the Hsp90 inhibitor Geldanamycin abrogated crosslinks to GR in all cases (Fig. 3a), strongly indicating that the observed intracellular FKBP51-Hsp90-GR complex depended on a functional Hsp90 machinery. Similar results were observed for FKBP52 (Fig. 3b). Next, we explored the effect of GR activation on selected FKBP crosslinks. Stimulation with the GR agonist Dexamethasone (Dex) induced rapid dissociation of the FKBP51-GR and FKBP52-GR complexes (Fig. 3c and d). The latter was surprising since FKBP52 has been thought to enter GR-Hsp90 complex after GR activation, possibly by replacing FKBP51 ^33^. Our findings unambiguously show that both FKBPs bind to GR in the apo form, prior to activation. Given the important role of helix α12 of GR in complexing FKBP51 (Fig. 2), we treated cells expressing a representative set of FKBP51 pBpa mutants with the GR antagonist Mifepristone (RU486), which is known to induce a GR conformation with a rearranged helix α12 ^34^. Like Dexamethasone, Mifepristone completely disrupted all FKBP51→GR and FKBP52→GR crosslinks (Extended Data Fig. 5a and b), indicating that the specific conformation of helix α12 is not important for dissociation of the FKBP-GR complexes in cells. The ligand sensitivity profile of the FKBP51-Hsp90-GR complex was confirmed by reverse crosslinks using selected GR mutants as proximity sensors (Extended Data Fig. 5c).

**Fig. 3.**
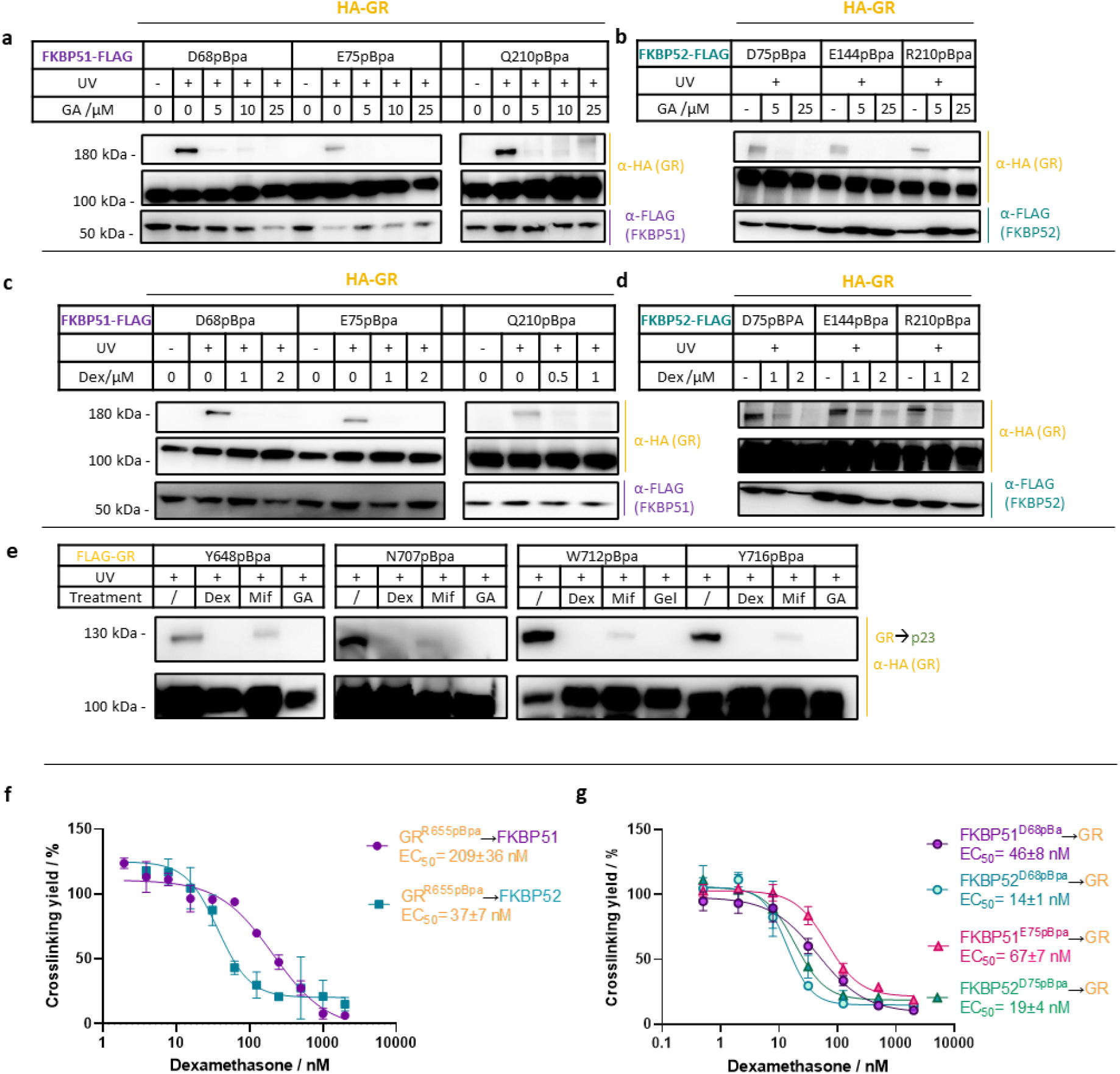
FKBP51, FKBP52, and p23 interact with the apoGR in a Hsp90-dependent manner. **a** + **b,** Hsp90 inhibitor (Geldanamycin, GA) treatment for 1 h disrupts FKBP51→GR (**a**) and FKBP52→GR crosslinks (**b**), suggesting Hsp90-dependent interactions. **c + d,** GR agonist (Dexamethasone, Dex) treatment for 1 h disrupts FKBP51→GR (**c**) and FKBP52→GR crosslinks (**d**), suggesting interactions with the apoGR under basal conditions. **e,** GR agonist (Dexamethasone, Dex, 5 μM) and Hsp90 inhibitor (Geldanamycin, GA, 5 μM) treatment for 1 h disrupts GR→p23 crosslinks; GR antagonist (Mifepristone, Mif, 5 μM) treatment for 1 h partially disrupts GR→p23 crosslinks. **f,** Dexamethasone dose-dependently disrupted the GR→FKBP51 and GR→FKBP52 interaction in cells as determined by ELISA after cellular stimulation for 30 min, in-cell photocrosslinking and cell lysis. **g,** Dexamethasone disrupts FKBP52→GR complexes more potently compared to FKBP51→GR complexes, as determined ELISA after cellular stimulation for 1 h, in-cell photocrosslinking and cell lysis. **f**+ **g,** For better comparison, the crosslinking yield is shown (mean±s.d.) with 0 μM Dex=100% crosslinking yield and –UV= 0%, plot represents data from biological replicates (*n*=3) per concentration. See Extended Data Fig. 5d–g for individual data points.

We also investigated ligand-sensitivity of GR crosslinks to p23. Stimulation with Geldanamycin, Dexamethasone or Mifepristone disrupted or severely compromised all GR→p23 crosslinks (Fig. 3e), suggesting that p23 too preferentially bound apoGR in a Hsp90-dependent manner.

The availability of intracellular GR activation-sensitive sensors allowed us to investigate the potency of GR agonists for GR activation in a subcomplex-resolved manner in cells. Using GR^R655pBpa^ co-expressed with FKBP51 or FKBP52 and ELISA as a readout, Dex titration revealed an apparent EC_50_ for GR activation of ~40 nM when starting from a GR complex containing FKBP52 (Fig. 3d). About 5-fold higher Dex concentrations were needed to activate GR from a FKBP51-containing GR complex. Similar EC_50_ values GR-FKBP51 complexes were obtained for two other GR mutants (Extended Data Fig. 5h).

The ELISA setup was also used to quantify the sensitivity of FKBP→GR crosslinks to GR activation, performed separately for FKBP51 and FKBP52. Using the same photocrosslinking positions for FKBP51 and FKBP52, FKBP52→GR crosslinks were found to be about 3–3.5-fold more sensitive to GR activation compared to FKBP51→GR crosslinks (Fig. 3g). While the absolute potencies for GR activation depended on GR expression levels, activation time, and photocrosslinking position, a similar potency ratio was found for matched FKBP52 vs FKBP51 mutants (Extended Data Fig. 5j and l). This shows that GR activation is facilitated in the context of a FKBP52-Hsp90-GR complex compared to GR in a FKBP51-Hsp90-GR complex, in line with the well-documented GR-facilitating effect of FKBP52 and the GR-repressing effect of FKBP51 ^35^.

## FKBP^FK1^ inhibitors reveal a multi-layered interaction with GR

The molecular basis for the differential effects of FKBP51 and FKBP52 on GR is still unknown. Several studies pointed towards a key role of the FK1 domains ^35–37^, in particular the proline-rich loop around L/P119 overhanging the FK506-binding site ^35^. However, studies on the role of the FK506-binding site itself have remained controversial ^27^. To investigate the role of the FKBP506-binding site for FKBP-Hsp90-GR complex assembly, we probed representative GR-photoreactive positions in FKBP51 for sensitivity towards the FKBP51 ligand SAFit2^38^. GR crosslinking was blunted for numerous positions in FKBP51^FK1^ (Fig. 4a, c and Extended Data Fig. 6a). Strikingly, however, none of the investigated positions in the FK2 or TPR domains of FKBP51 were affected (Fig. 4b, c and Extended Data Fig. 6a). Similar results were obtained using the bicyclic FKBP ligand 18^(S)-Me^ (Ref 39) (Extended Data Fig. 6b) and when probing FKBP52 (Extended Data Fig. 6c and d). SAFit2 and the bicyclic FKBP ligand disrupted crosslinks in FKBP51^FK1^ and FKBP52^FK1^ at concentrations consistent with their potency to occupy intracellular FKBP51 or FKBP52 ^40^ (Fig. 4d). This identifies the FK2 and TPR domains as the major drivers for the assembly of FKBP51- or FKBP52-Hsp90-GR complexes in cells, while many of the FK1 contacts are dispensable.

**Fig. 4.**
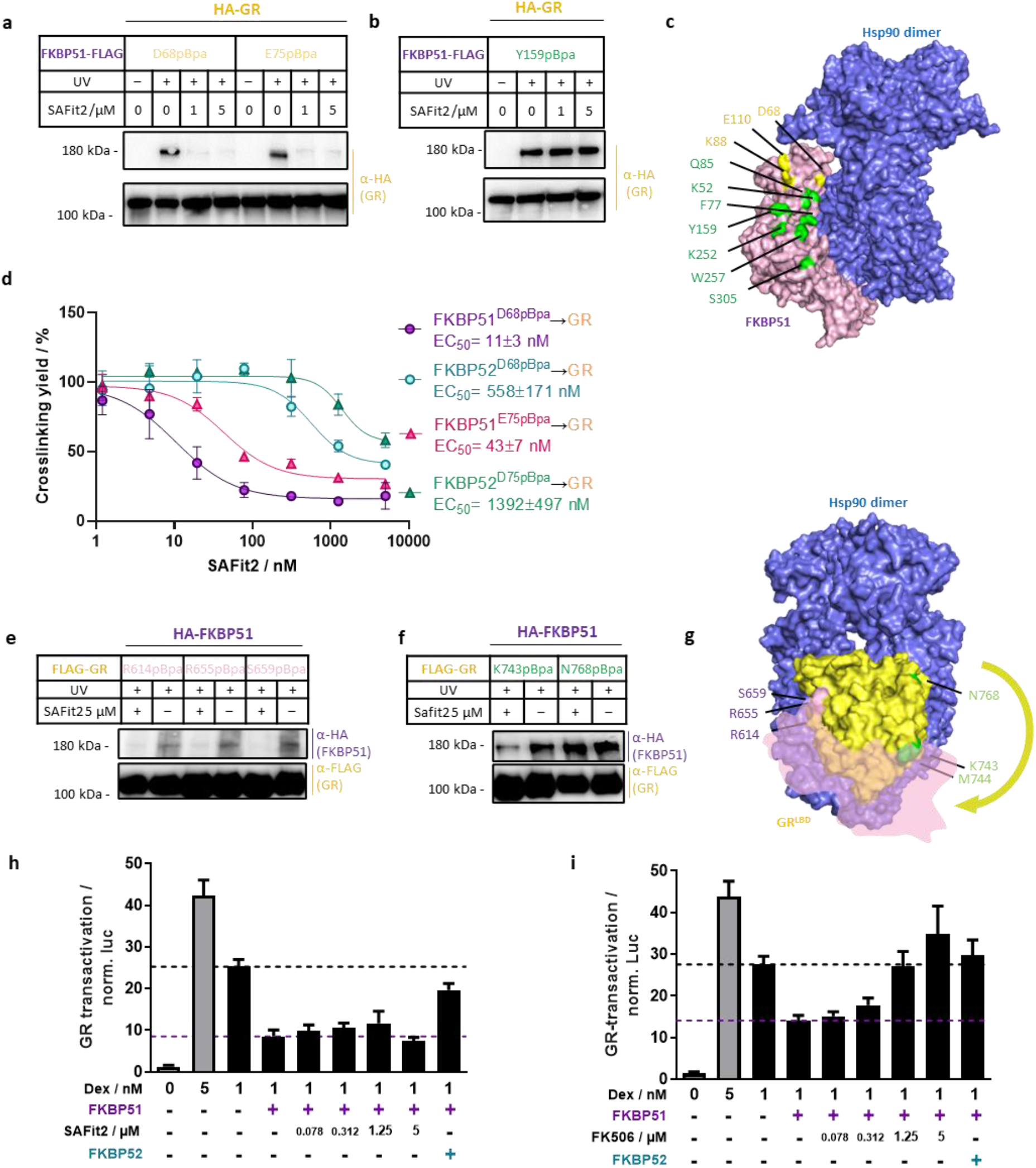
Inhibition of FKBPs only partially disrupts FKBP-GR interactions. **a–c,** Treatment with the FKBP51 inhibitor (SAFit2) for 1 h disrupts FKBP51→GR crosslinks in the FK1 (**a**) but not the FK2 (**b**) and TPR domains. **a** + **b,** Western blots of exemplary FKBP51 pBpa mutants expressed and photocrosslinked in HEK293 cells co-overexpressing HA-tagged GR. UV light-induced HA-reactive bands at a size of approx. 180 kDa are indicative of the mutated position being in proximity to GR. **c,** FKBP51→GR crosslinks tested for SAFit2 sensitivity were mapped to the structure of the FKBP51-Hsp90 complex (PDB: 7J7I, p23 omitted for clarity). SAFit2-sensitive crosslinks are shown in yellow, SAFit2-insensitive crosslinks (i.e., persisting in the presence of SAFit2) are shown in green. **d,** SAFit2 dose-dependently disrupts photocrosslinking positions in FKBP51^FK1^ and FKBP52^FK1^, determined by ELISA. For better comparison, the crosslinking yield is shown (mean±s.d.) with 0 μM Dex=100% crosslinking yield and –UV= 0%, plot represents data from biological replicates (*n*=3) per concentration. See Extended Data Fig. 6f + g for individual data points. **e** + **f,** SAFit2-sensitivity (**e**) or resistance (**f**) of exemplary GR pBpa mutants, detected by Western blots after expression and photocrosslinked in HEK293 cells with co-overexpressing HA-tagged FKBP51. **g,** GR→FKBP51 crosslinks tested for SAFit2 sensitivity were mapped to the structure of the GR-Hsp90 complex (PDB: 7KRJ, p23 omitted for clarity). SAFit2-sensitive crosslinks are shown in pink, SAFit2-insensitive crosslinks (i.e., persisting in the presence of SAFit2) are shown in green. **h** + **i,** GR transactivation was measured by reporter gene assays (mean±s.d.) in HEK293 cells transiently co-transfected with the dual reporters pGL4.36 (MMTV-luc2p), pGL4.74 (TK-hRluc) as well as expression plasmids for human GR and optionally FKBP51 and/or a 3-fold excess of FKBP52. See Extended Data Fig. 7a + b for individual data points. Treatment with FK506 (**i**) but not SAFit2 (**h**) dose-dependently blocks FKBP51-induced GR suppression. Data from biological replicates (*n*=6) per concentration.

We further probed representative FKBP51-photoreactive positions in GR for SAFit2 sensitivity. Notably, positions clustering around R614, R655 and E659 were found to be SAFit2-sensitive (Fig. 4e, g and Extended Data Fig. 6e), whereas crosslinks clustered around helix α12 were not affected (Fig. 4f, g and Extended Data Fig. 6e). Together with the ligand sensitivity findings for FKBP51 (Fig. 4a and b), this suggests that residues around GR α-helix 12 form an interface with the FK2 and/or TPR domain of FKBP51, while the region around the tip of α5 and the α7-α8 loop of GR contact FKBP51^FK1^, consistent with the rotation postulated for GR above.

To explore the functional consequences of ligand-induced FK1 remodelling, we performed GR reporter gene assays in HEK293 cells. As previously reported ^37^, co-overexpression of FKBP51 reduced GR transactivation. However, the GR-repressing effect of FKBP51 was not affected by SAFit2 treatment (Fig. 4h), up to a dose >100-fold higher than necessary for FKBP51^FK1^ remodelling (Fig. 4d). The GR-repressing effect of FKBP51 was principally reversible, as shown by co-overexpression of FKBP52. Similar findings were observed for the bicyclic FKBP ligand 18^(S)-Me^ (Extended Data Fig. 7c). However, the larger macrocyclic ligand FK506 ^41^, which protrudes from the FK506-binding site of FKBP51 much more than SAFit2 ^42^ or 18^(S)-Me 39^ (Extended Data Fig. 7c), reverted FKBP51-induced GR-suppression at doses consistent with intracellular FKBP51 occupancy ^40^ (Fig. 4i).

Collectively, these findings show that the FK506-binding site itself is not required for the GR-suppressing effect of FKBP51 but that larger FKBP ligands can perturb functionally relevant interactions.

## Discussion

Steroid hormone receptor functioning relies crucially on the Hsp90 chaperone machinery for loading and activation by hydrophobic steroids. The minimal chaperone machinery, best defined for GR, consists of five factors (Hsp90, Hsp70, Hsp40, Hop, and p23) ^28^. Two key intermediate states of the basic GR chaperoning cycle have recently been structurally elucidated by the Agard group ^22,23^ (Fig. 5), which represent early chaperoning and late activated states, respectively.

**Fig. 5.**
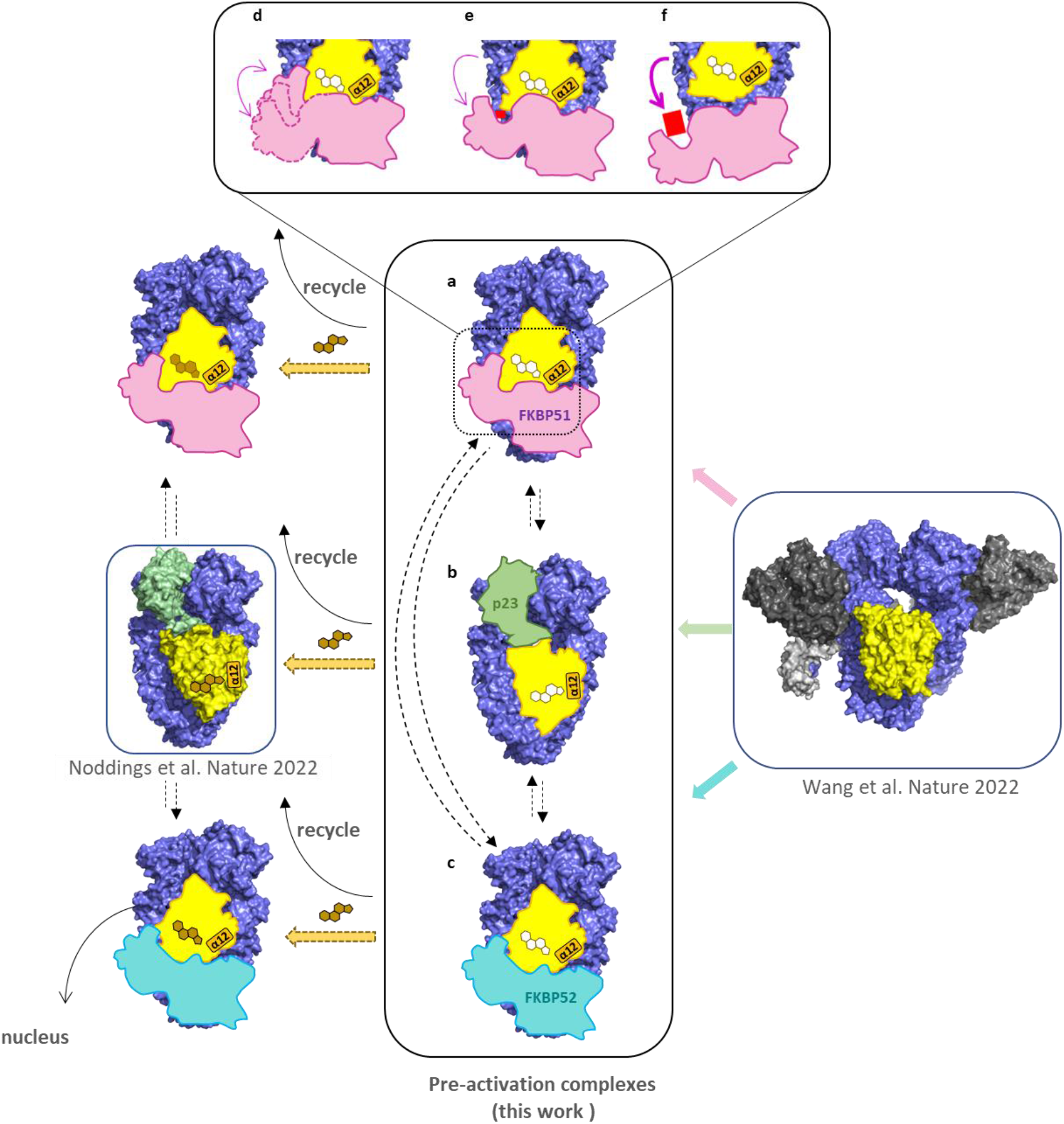
Co-chaperones preferentially engage apoGR in pre-activation complexes in human cells. After Hop-mediated transfer from Hsp70 to Hsp90 (right), apoGR transitions further to FKBP51 (**a**), p23 (**b**), or FKBP52 (**c**) containing pre-activation complexes, which represent the predominantly populated FKBP/p23-GR species in cells. The pre-activation complexes can progress by GR ligand activation to Hsp90-cochaperone complexes containing activated GR (right) or recycle by disassembly of the heterocomplexes. **d,** Loops adjacent to the FK506-binding pocket are flexible and contact the GR to regulate GR ligand binding. **e,** smaller FKBP ligands stabilize the loops, disrupt some GR-contacts of the FK1 domain, but spare the GR-regulatory contacts and the association with the FK2 and TPR domains. **f,** Larger FKBP ligands disrupt GR-regulatory contacts.

Higher eukaryotes have evolved FKBP51 and FKBP52 as additional Hsp90 co-chaperones to super-regulate steroid receptor signaling, with essential physiological roles in mammals ^32^. While FKBP52s main function seems to be to support optimal receptor activation (incl. subsequent transport to the nucleus ^37,43^), FKBP51 is a GR repressor and key effector of an ultrashort negative feedback loop ^19^. However, how and precisely when FKBPs act on GR in the chaperone cycle has been matter of debate.

By using photocrosslinking in intact cells we capture FKBP51 and FKBP52 in their functionally relevant states, which have so far eluded any purification or reconstitution attempts. The studied system recapitulates the hallmarks of the FKBP-Hsp90-GR machinery, incl. Hsp90 dependence and facilitated GR activation by FKBP52 compared to FKBP51. Our results show that in the absence of hormone FKBP51, FKBP52 as well as p23 predominantly reside in apoGR-Hsp90 containing complexes in human cells (Fig. 5a–c). These states can either progress towards activation by GR ligand stimulation or recycle through the Hsp90 machinery. Both of these processes are fast (<10min, Extended Data Fig. 5l) and complete. We show that either pathway leads to complete disassembly of the entire complex, since none of the investigated interactions survived as long as hormone stimulation persists or reassembly, respectively, is blocked.

Our findings do not exclude the existence of FKBP-Hsp90 complexes with activated, ligand-bound GR, which were previously suggested especially for FKBP52 ^33,44^. However, these complexes are likely short-lived in cells and quickly disassemble after the (co)-chaperone machinery has done its job to deliver activated GR.

Our results also provide a picture for the overall architecture of the relevant FKBP/p23-Hsp90-apoGR pre-activation complexes (Fig 5). The p23-Hsp90-apoGR complex resembles the maturation complex (p23-Hsp90-GR^Dex-bound^) ^23^ elucidated by cryo-EM. For FKBP-containing complexes, the GR has to rearrange, e.g., by rotation of the GR^LBD^, which stays tethered to the client-binding channel of Hsp90 via pre-helix α1 of GR. This rotation will abrogate p23-GR contacts observed in the absence of FKBPs (Fig. 2e and Ref 23), which may be partially replaced by contacts with the tip of FKBP51. FKBP51, FKBP52, and p23 are likely inter-exchangeable in the Hsp90-apoGR complex, either directly at preassembled Hsp90-apoGR complexes or via Hsp90/Hsp70/Hsp40/Hop-mediated dis- and reassembly.

Our results confirm the ligand binding domain of GR as the predominant interaction site for FKBPs, in line with prior studies ^36^, and clearly show that FKBP51 and FKBP52 engage in extensive contacts with GR^LBD^. The identified FKBP→GR interaction surface is largely consistent with the conformation of FKBP51 bound to Hsp90 in the absence of GR ^29^ and requires only minor rearrangements of the FK1 and FK2 domains. The GR has to reorient more profoundly compared to available structures ^22,23^ to allow for the observed interaction surface. FKBP51 wraps around the GR^LBD^ from helix α12 to the α7-α8 loop, covering the α1-α3 loop, where mutations were previously shown enhance GR repression by FKBP51 ^45^.

A key question is how FKBP51 and FKBP52 exert strikingly opposing effects on GR signaling in spite of high structural and sequence similarity. In line with their similarities, most of the GR-interacting interface is identical between FKBP51 and FKBP52. While the inter-domain arrangements of FKBP52 were shown to be flexible ^21^, it is now clear that FKBP51 and FKBP52 adopt an almost identical conformation in complex with Hsp90-apoGR. Previous domain swapping analyses pointed to the FK1 domain as a major factor underlying the opposing effects of FKBP51 and FKBP52. Indeed, the most diverging GR interaction pattern was located around the FK506-binding site (Fig. 1b and d), especially in the proline-rich loop overhanging this site (P109-P124, Extended Data Fig. 8). Intriguingly, swapping L119 in FKBP51 for P119 as in FKBP52 was previously shown to impart GR-stimulating activity to FKBP51 ^35^. Crystal structures previously showed the proline-rich loop to be flexible in FKBP51 ^30,38,46^ as well as in FKBP52 ^38,47^, and NMR studies suggested differential dynamics of this loop as well as the beta bulge on the opposite side of the FK506-binding pocket to account for the diverging effects of the large FKBPs for GR regulation ^48^.

The precise role of FKBP51^FK1^ and FKBP52^FK1^ has important implications for FKBP-directed drug discovery but previous pharmacological studies have been inconclusive ^44,49,50^. We show that compounds binding to the FK506-binding site of FKBP51 or FKBP52 clearly remodel the contacts between the FK1 domain and GR, while sparing the docking of the GR to the FK2 and TPR domains. For smaller ligands, the remaining FK1 domain contacts are sufficient to maintain the GR-suppressive effect of FKBP51 (Fig. 5e). Larger ligands, however, are more disruptive and abolish the GR-suppressive effect of FKBP51 (Fig. 5f).

Our proposed binding modes for the FKBP-Hsp90-apoGR preactivation complexes are remarkable consistent with two structures for the related complexes containing Dex-bound GR, recently obtained by the Agard group by cryoEM after *in vitro* reconstitution (Noddings *et al*., submitted). Shared features are the intensive contacts of all three FKBP domains with GR^LBD^, the highly similar interaction pattern for FKBP51 and FKBP52, the rotation of GR^LBD^ compared to the p23-Hsp90-GR^LBD+Dex^ complex, the stronger, well defined docking of the FK2 and TPR domains to GR^LBD^ compared to a more dynamic, partially displaceable FK1-GR^LBD^ interaction, and conclusions regarding the role of the FK506-binding sites. The observed interdomain assembly seems to be a strong scaffold that can accommodate GR in pre- and post-activation states.

Systematic site-specific incorporation of photoreactive unnatural amino acids has been used before to probe interactions of small or peptide ligands with receptors in mammalian cells ^51–54^ or between proteins in yeast, *E.coli* or *in vitro* ^55,56^. However, to the best of our knowledge this is the first example, where the architecture of a multi-protein complex has been determined directly in intact mammalian cells. Notably, this enables determination of apoGR-containing complexes that so far have eluded any biochemical purification of reconstitution attempts. The streamlining of large-scale photocrosslinking screening (>600 tested positions) allowed to reconstruct the overall architecture of metastable complexes that have eluded structural or tailored functional studies so far. Moreover, the use of proximity sensors enabled to trace the remodelling of complexes at much higher detail than previously possible in cells. Systematic surface mapping might be a general approach for low-resolution in-cell structural biology that is poised to interface with functional cellular studies.

Taken together, we capture FKBP51 and FKBP52 in the states of GR suppression and pre-activation, respectively, thought to be crucial for stress reactivity regulation in humans and for sexual development in mammals. The proposed interaction modes are extendable to other FKBP-regulated steroid hormone receptors such as progesterone, estrogen or androgen receptors, with important implications for breast and prostate cancers ^13,14^. We define the overall architecture of the FKBP-apoGR complexes and provide the rationale for developing FKBP-directed ligands with higher levels of efficacy.

## Methods

### Golden Gate reaction

All Golden Gate reactions were performed as describe in Püllmann et al.^1^ PCR reactions were carried out using standardized conditions in a 96-well format to generate a fragment 1 and a fragment 2 for each mutant. The final Golden Gate reaction volume contained 1× concentrated T4 ligase buffer (NEB), 250 ng Golden Gate vector (pcDNA3-based carrying a twin Strep-FLAG tag, synthesized by BioCat, Heidelberg, Germany) 2 U BSA HFv2 (NEB), 1 U T4 ligase (NEB) and the fragment 1 & 2 for each mutant. Golden Gate reactions were carried out in a 96-well plate (30 cycles of 3 min 37°C (enzymatic digest) and 3 min 16°C (ligation reaction), followed by a final restriction step 3 min 37°C before the heat inactivation 20 min 80°C), followed by transformation into chemocompetent *E. coli* DH5α cells (NEB) by heat shock procedure. Selected mutants were sequenced to ensure the side specific mutation.

### Amber suppression

Amber suppression was carried out using a modified protocol of Serfling & Coin ^2^. Cell culture experiments were performed with Human embryonic kidney (HEK) 293 cells that were cultured in Dulbecco’s modified Eagle’s medium (DMEM) with 10% fetal bovine serum (FBS) and 1% penicillin-streptomycin solution at 37°C with 5% CO2. Cells were grown overnight to 50% confluence. Transfection was performed according to Lipofectamin 2000 (Invitrogen) cell transfection protocol. The transfection mix contained the pBpa tRNA synthetase p_NEU_EBpaRS_4xBstYam and four copies of U6-BstYam expression cassettes ^2^ (kind gift of Irene Coin, Leipzig) and the corresponding TAG mutant of either FKBP51, FKBP52, or GR. pBpa (Thermo Scientific) was added to the media at a concentration of 500 μM. The cells were incubated for 42 h to allow for expression of pBpa-containing proteins, followed by optional pharmacological treatment (as indicated in the figures), and UV-irradiated for 30 min on ice.

### Western blot

Cells were lysed in NETN buffer (100 mM NaCl, 20 mM Tris pH 8, 0.5 mM EDTA, 0.5 % Nonidet P-40, protease inhibitor cocktail (Roche)). The proteins were separated by SDS-PAGE and transferred to a nitrocellulose membrane (Amersham). The protein on the membrane were probed with antibodies (either anti-HA (Roche),-FLAG (Sigma), or -p23 (SCBT)) and detected with Immobilon Western (Millipore). The band intensity was measured with an image analyzer (Fuji Photo Film).

### ELISA

ELISA plates (Maxisorp, 96 well plate, Invitrogen) were coated with 100 μL of 7.5 ng/μL Streptactin (IBA) in PBS and the plates were incubated at 4°C overnight. Plates were washed with PBS/0.05% Tween20 (ELISA washer: BioTek ELx405 Select CW) and blocked with 5% milk powder in TBS at 4°C overnight. Plates were washed with PBS/0.05% Tween20 and plates were stored at 4°C. For ELISA, 70 μL PBS was added to each well. 30 μL of cell lysate were added and the plate was incubated at 4° C overnight. ELISA plates were washed with PBS/0.05% Tween20. Then, the antibody solution (1:5000 anti-FLAG in 5% BSA in 1×TBST or 1:1000 anti-HA in PBS) were added and the plate was incubated for 2 h at room temperature. Plates were again washed and the TMB substrate (ThermoFisher) were added. After 10 min incubation in the dark, the enzymatic reaction was quenched by adding 0.18 M H_2_SO_4_. Absorption at 450 nm was measured (Tecan Spark). Data were analysed with GraphPad Prism 9.

### Reporter Gene Assays

Prior to transfection HEK293 cells were seeded at a density of 1×10^4^ per well in poly-D-lysine coated 96 well plates. After overnight attachment the dual reporter plasmids (pGL4.36, pGL4.74 (Promega)), prK5-HA-GR as well as prK5-FKBP51-Flag and prK5-FKBP52-Flag were introduced by polyethyleneimine-mediated transient transfection. On the next day, the cells were stimulated with Dexamethasone and co-treated with FKBP ligands or DMSO for 24 hours. Subsequently, the cells were washed with DBPS and lysed in 60 μL Passive lysis buffer (Promega, E1910). For measurement, 20 μL cell lysates were transferred to white 96 half area plates (Greiner, 675075) and reporter expression was quantified using a Tecan Spark and Dual-Glo^®^ Luciferase Assay System (Promega, E2920) according to a manufacturers instruction.

## Acknowledgements

We thank Martin Weissenborn, Pascal Püllmann, and Chris Ulpinnis (University of Halle) for suggestions and training on the golden gate mutagenesis protocol, Irene Coin (University of Leipzig) for plasmids and suggestions for pBpa incorporation in mammalian cells, and Jürgen Kolos and Tim Heymann for samples of 18^(S)-Me^ and SAFit2, respectively. We are indebted to Malin Wilfinger and Jan-Philip Kahl for the cloning GR mutants and helping to establish the ELISA and to Francois Halloy for preliminary work on reporter gene assays. This work was supported by funding from the HMWK (LOEWE-Schwerpunkt TRABITA) and the BMBF (16GW0290K) to F.H.

## Author Contributions

A.B & S.E. designed and executed all photocrosslinking experiments and subsequent analysis. T.M.G performed and analyzed the reporter gene assays. F.H. conceived the project. All authors interpreted the results, A.B. and F.H. wrote the manuscript, approved by all authors.

## Additional Information

Correspondence to Felix Hausch

